# Retinal ganglion cells harboring the OPTN(E50K) mutation exhibit neurodegenerative phenotypes when derived from hPSC-derived three dimensional retinal organoids

**DOI:** 10.1101/820159

**Authors:** KB VanderWall, KC Huang, Y Pan, SS Lavekar, CM Fligor, A Allsop, K Lentsch, P Dang, C Zhang, HC Tseng, TR Cummins, JS Meyer

## Abstract

Retinal ganglion cells (RGCs) serve as the primary connection between the eye and the brain, with this connection disrupted in glaucoma. Numerous cellular mechanisms have been associated with glaucomatous neurodegeneration, and useful models of glaucoma allow for the precise analysis of degenerative phenotypes. Human pluripotent stem cells (hPSCs) serve as powerful tools for studying human neurodegenerative diseases, particularly cellular mechanisms underlying degeneration. Thus, efforts were initially focused upon the use of hPSCs with an E50K mutation in the Optineurin (OPTN) gene. CRISPR/Cas9 gene editing was used to introduce the OPTN(E50K) mutation into existing lines of hPSCs, as well as the generation of isogenic control lines from OPTN(E50K) patient-derived hPSC lines. OPTN(E50K) RGCs exhibited numerous neurodegenerative deficits, including neurite retraction, autophagy dysfunction, apoptosis, and increased excitability. The results of this study provide an extensive analysis of the OPTN(E50K) mutation in hPSC-derived RGCs, with the opportunity to develop novel treatments for glaucoma.

## Introduction

Glaucoma is a devastating optic neuropathy which causes the progressive degeneration of retinal ganglion cells (RGCs), leading to irreversible loss of vision and eventual blindness (Quigley, 2011). Various animal models have been developed that have led to a greater understanding of glaucomatous neurodegeneration(Agostinone and Di Polo, 2015; Cueva Vargas et al., 2015; Kalesnykas et al., 2012; Williams et al., 2013), although these models often exhibit physiological features that do not precisely reflect those present within human patients. Furthermore, recent studies demonstrating significant variability between rodent and primate RGCs (Peng et al., 2019) suggest that there may be important differences in how these cells respond to glaucomatous injuries between species. As such, there is a strong need to develop new approaches to complement these glaucoma models in order to determine RGC pathogenesis and mechanisms leading to their degeneration and death.

Human pluripotent stem cells (hPSCs) provide an attractive option as a model for studies of cellular development and disease progression as they can be cultured indefinitely and induced to differentiate into all cell types of the body (Thomson et al., 1998), including RGCs (Deng et al., 2016; Gill et al., 2016; Langer et al., 2018; Ohlemacher et al., 2016; Riazifar et al., 2014; Tanaka et al., 2015; Teotia et al., 2017a). When harboring genetic mutations associated with disease states, the derivation of these cells from patient-specific sources allows for the ability to study mechanisms underlying diseases such as glaucoma (Inagaki et al., 2018; Minegishi et al., 2013; Ohlemacher et al., 2016; Teotia et al., 2017b). Additionally, gene editing approaches including CRISPR/Cas9 technology allow for the ability to create isogenic controls from these patient-derived cells and also introduce disease-causing mutations in unaffected cells, leading to the generation of new disease models (Cong et al., 2013). Among the gene mutations associated with glaucoma, those mutations in the Optineurin (OPTN) gene are known to result in glaucomatous neurodegeneration in the absence of elevated intraocular pressure (Minegishi et al., 2016). These mutations directly affect retinal ganglion cells, with the OPTN(E50K) mutation previously shown to result in a particularly severe degenerative phenotype (Chalasani et al., 2014; Inagaki et al., 2018; Meng et al., 2012; Minegishi et al., 2013; Ohlemacher et al., 2016; Tseng et al., 2015; Ying et al., 2015). As such, the generation of hPSCs harboring the OPTN(E50K) mutation, along with corresponding isogenic controls, allows for the in vitro analysis of mechanisms underlying the degeneration of RGCs in glaucoma.

Thus, the efforts of the current study were initially focused upon the generation of hPSCs with the OPTN(E50K) mutation as well as their corresponding isogenic controls. These models were generated through the use of CRISPR/Cas9 gene editing allowing for the generation of isogenic controls. Upon initial differentiation of these cells into three-dimensional retinal organoids and subsequent RGC purification, OPTN(E50K) and isogenic control cells both developed in a similar manner, yielding a comparable number of RGCs. Conversely, at later stages of RGC maturation, OPTN(E50K) RGCs demonstrated numerous characteristics associated with glaucomatous neurodegeneration, including neurite retraction, autophagy dysfunction, and increased excitability. The results of this study provide the most comprehensive analysis of glaucomatous neurodegeneration to date using hPSC-based models, including the demonstration of the use of CRISPR/Cas9 gene editing to provide essential disease models and corresponding isogenic controls for glaucoma. The identification of numerous neurodegenerative phenotypes associated with the OPTN(E50K) glaucoma state allow for the exploration of therapeutic intervention including pharmacological screening and cell replacement therapies.

## Results

### CRISPR/Cas9-Edited OPTN(E50K) Disease Models and Isogenic Controls

The ability to precisely edit genes in hPSCs through CRISPR/Cas9 gene editing technology allows for new insights into disease modeling including the generation of isogenic controls (Cong et al., 2013). When properly applied, CRISPR/Cas9 gene editing allows for the distinction between cell line variability and mutation-causing disease phenotypes. As such, CRISPR/Cas9 gene editing was utilized in the current study to examine the E50K mutation in the Optineurin protein, a known genetic determinant for normal tension glaucoma.

In order to introduce the OPTN(E50K) mutation in hPSCs, the homology directed repair (HDR) template was designed to include the mutant nucleotide c.148G>A that altered the 50^th^ amino acid from glutamic acid (E) to lysine (K), with an additional two silent mutations (c.144G>C and c.150G>A). The first silent mutation altered a Hpy188III restriction site, which was used for PCR screening of prospective gene-edited clones. The second silent mutation was introduced to prevent Cas9 from cutting the donor template by modifying the PAM site (Figure 1). A plasmid containing the OPTN(E50K) HDR template as well as the guide RNA (gRNA) was co-transfected with pCas9-GFP, with the GFP signal used to sort for potentially edited cells. Clonal populations were screened by PCR amplification of the edited region and enzymatic digestion with Hpy188III, with prospectively edited clones further sequenced to ensure proper editing of the target gene. Likewise, patient-specific hiPSCs harboring the OPTN(E50K) mutation utilized a similar strategy to correct the OPTN(E50K) mutation. The results of these experiments demonstrated the successful establishment of hPSC lines harboring the OPTN(E50K) mutation along with their corresponding isogenic controls.

**Figure 1:**
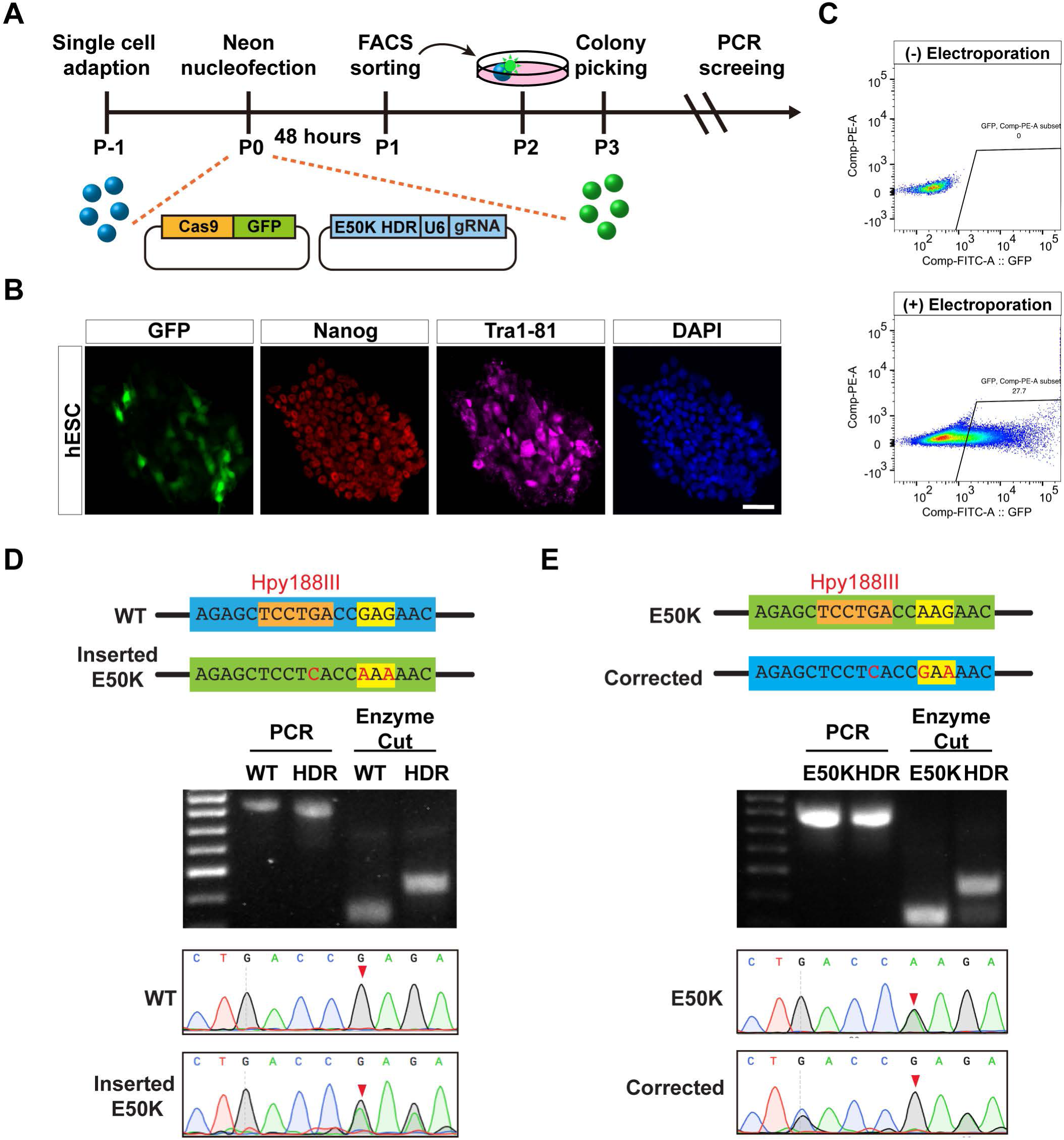
CRISPR/Cas9 gene editing for the generation of OPTN(E50K) hPSCs and corresponding isogenic controls. (a) A schematic displays the work flow for the CRISPR/Cas9 gene editing of hPSCs. (b) Fluorescence microscopy demonstrated a subset of GFP-positive cells in hPSCs expressing Nanog and Tra1-81 after electroporation. (c) Flow cytometry graphs indicated the gating of GFP positive cells after 48 hours of electroporation. (d-e) CRISPR/Cas9 gene edited cells were screened for the insertion of OPTN(E50K) mutation in wild-type hPSCs (d) and correction of the mutation in patient-derived hPSCs (e) by restriction digestion with Hpy188III and Sanger sequencing. The 50^th^ codon of the OPTN gene is highlighted in yellow. The red arrows highlight the insertion or correction of the OPTN(E50K) missense mutation site in Sanger sequencing. Scale bar equals 50μm in b.

### OPTN(E50K) glaucomatous RGCs demonstrate morphological deficits and gene downregulation after prolonged culture

The degeneration of RGCs in glaucoma severely affects the visual pathway, leading to blindness (Quigley, 2011). This degeneration is commonly associated with advanced age, as the initial development of the retina is unaffected by the disease state. As such, the early differentiation of OPTN(E50K) and isogenic control hPSCs yielded similar formation of optic vesicle-like and optic cup retinal organoids and a comparable number of RGCs (Supplemental Figure 1). Previous studies in animal models of RGC damage have demonstrated significant neurite retraction and dendritic remodeling in response to RGC injury and disease (Agostinone et al., 2018; Agostinone and Di Polo, 2015; Kalesnykas et al., 2012; Williams et al., 2013). To examine if this phenomenon could be recapitulated in an hPSC model, OPTN(E50K) and isogenic control RGCs were examined in a temporal fashion for their morphological characteristics associated with neuronal maturation (Figure 2). Representative inverted fluorescent images and neurite tracings of OPTN(E50K) and isogenic control RGCs revealed high degrees of similarity in neuronal maturation from 1 to 3 weeks after purification. By 4 weeks of maturation, however, OPTN(E50K) RGCs displayed deficits in neurite complexity, cell body size, neurite length and expression of various pre- and post-synaptic proteins (Supplementary Figure 2). These results indicated that OPTN(E50K) and isogenic control RGCs matured similarly during the early stages of neuronal growth in regards to their morphology but after prolonged culture, OPTN(E50K) RGCs significantly reduced their morphological complexity, similar to what has been previously observed for glaucomatous RGCs in vivo.

**Figure 2:**
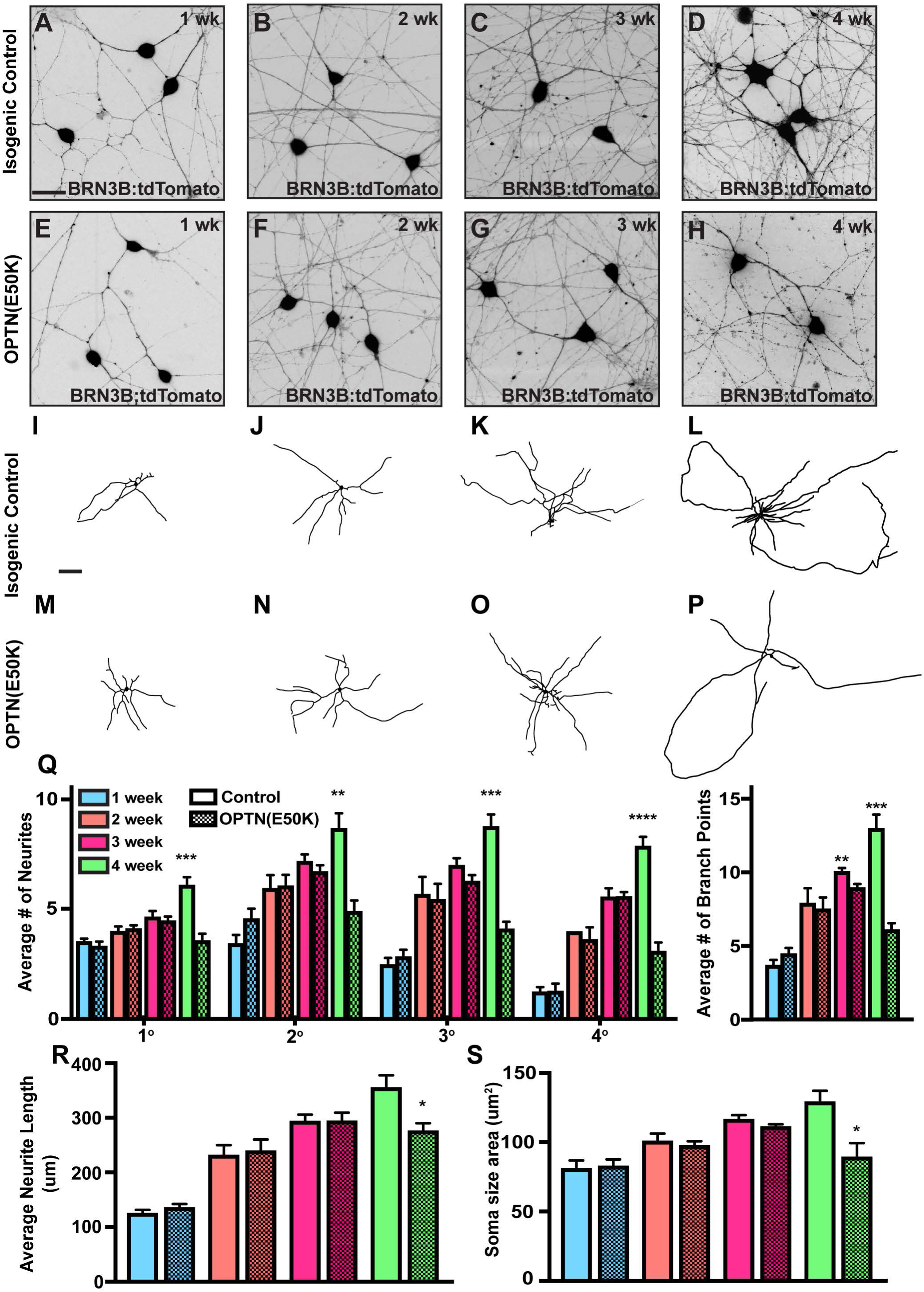
OPTN(E50K) RGCs demonstrate neurite deficits at later stages of maturation. (a-h) Fluorescent images and (i-p) representative neurite tracings of isogenic control and OPTN(E50K) RGCs from 1 to 4 weeks after purification displayed similar neurite complexity during early timepoints, but significant neurite retraction at 4 weeks in OPTN(E50K) RGCs. (q) Quantification for neurite complexity revealed a significantly lower number of ordered neurites and branch points in OPTN(E50K) RGCs after 4 weeks. (r-s) OPTN(E50K) RGCs also demonstrated significantly shorter neurites and smaller cell soma sizes compared to isogenic controls by 4 weeks of maturation. Error bars represent SEM (n=4 biological replicates using H7 and H7-OPTN(E50K) hPSCs). (*p < 0.05; **p < 0.01; ***p < 0.001, ****p < 0.0001). Scale bars equal 25μm for a-h and 100μm for i-p.

Previous studies in animal models of RGC injury have also demonstrated the down-regulation of RGC-associated proteins prior to the degeneration of these cells (Soto et al., 2008; Weishaupt et al., 2005). As such, efforts were made to determine if RGC-associated proteins were similarly downregulated in OPTN(E50K) RGCs compared to isogenic controls (Figure 3). As a percentage of the total cell population, the expression of the RGC transcription factors BRN3B and ISLET1 were significantly decreased in OPTN(E50K) RGCs compared to isogenic controls. To determine if this decrease was a result of gene downregulation or loss of cells, the expression of each was compared to the expression of the RGC-associated cytoskeletal marker MAP2. No significant differences were observed in the percentage of MAP2-positive cells between OPTN(E50K) and isogenic control RGCs, however the colocalization of either BRN3B or ISLET1 with MAP2 was significantly reduced in the OPTN(E50K) condition. These results suggested an early downregulation of RGC-associated transcription factors in response to the OPTN(E50K) mutation, indicative of an increased susceptibility to subsequent glaucomatous neurodegeneration.

**Figure 3:**
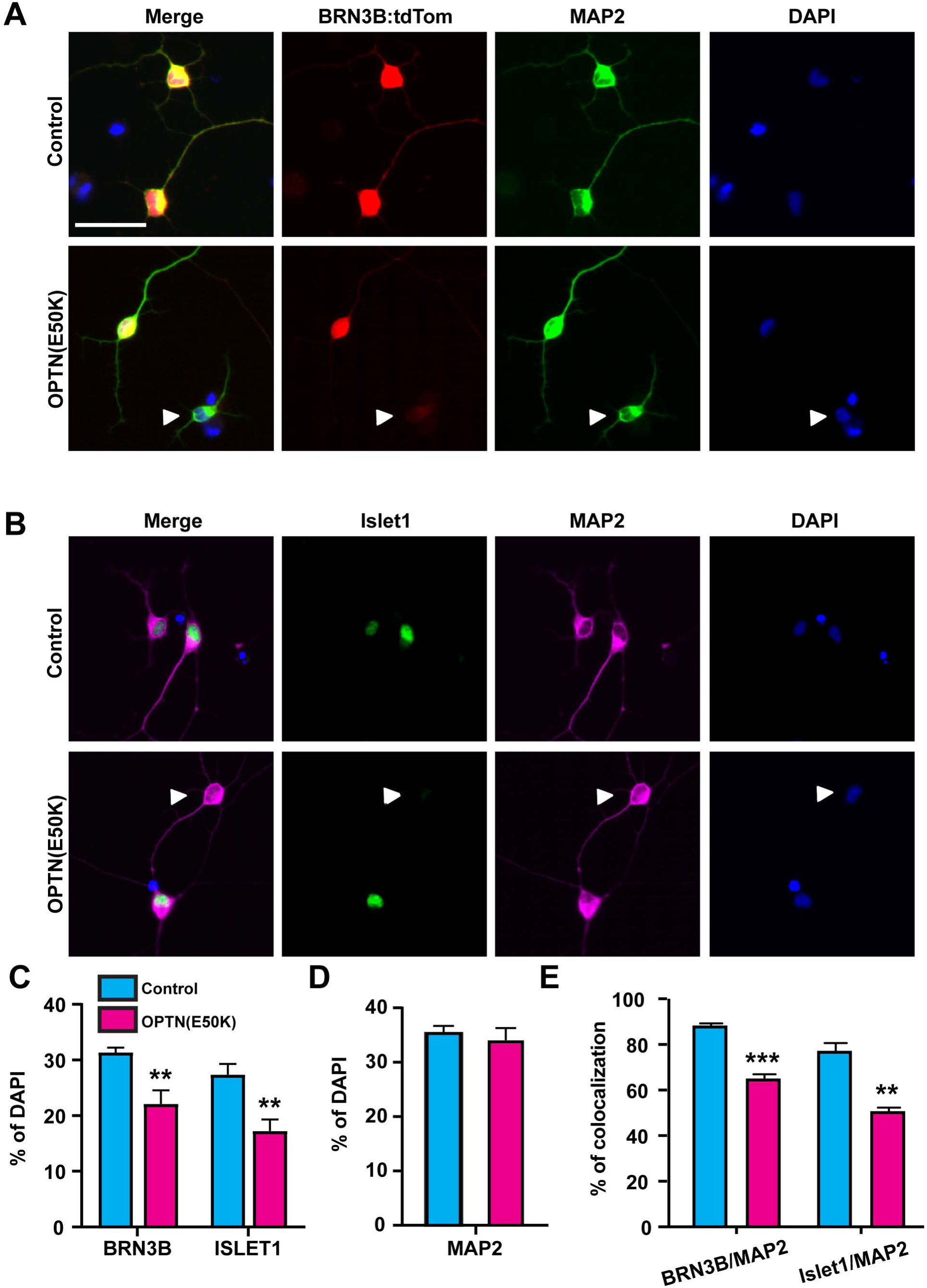
OPTN(E50K) RGCs downregulate RGC-associated transcription factors. (a-b) Immunostaining displayed the expression of RGC-associated transcription factors BRN3B:tdTOMATO and ISL1 correlated with the RGC-associated cytoskeletal marker MAP2. (c) Quantification of immunostaining results demonstrated a significant decrease in the expression of BRN3B and ISL1 in OPTN(E50K) RGCs relative to the total number of DAPI-positive cells. (d) No significant changes were observed in the expression of MAP2 between conditions. (e) Lastly, the co-localization of BRN3 and ISL1 with MAP2 was significantly decreased in OPTN(E50K) RGCs compared to isogenic controls. Error bars represent SEM (n=3 biological replicates using OPTN(E50K) hiPSCs, OPTN- corrected hiPSCs, H7 and H7(E50K) hPSCs). **p < 0.01; ***p < 0.001). Scale bar equals 25 μm for both a and b.

### Functional consequences of the OPTN(E50K) glaucomatous mutation

RGCs must be functionally active in order to transmit visual information, with propagation of this information achieved through the excitability of these cells and their firing of action potentials (Wang et al., 1997). Previous studies have demonstrated changes in RGC excitability associated with the glaucomatous disease state (Cueva Vargas et al., 2015) and as such, patch clamp analyses were utilized to determine if functional changes were present in OPTN(E50K) RGCs compared to isogenic controls (Figure 4). Within 4 weeks of RGC maturation, both OPTN(E50K) and isogenic control RGCs demonstrated functional properties. No significant changes were detected in the resting membrane potential, although OPTN(E50K) RGCs displayed a significantly lower cell capacitance and action potential current threshold as well as higher input resistance compared to isogenic control cells, suggesting the possibility for changes to their excitable properties. To test this, current clamp recordings were performed and upon injection of small amounts of current, OPTN(E50K) RGCs fired significantly more action potentials compared isogenic controls. These results demonstrated an increased excitability of OPTN(E50K) RGCs, suggesting that excitotoxicity may play a key role in the early stages of RGC degeneration in glaucoma.

**Figure 4:**
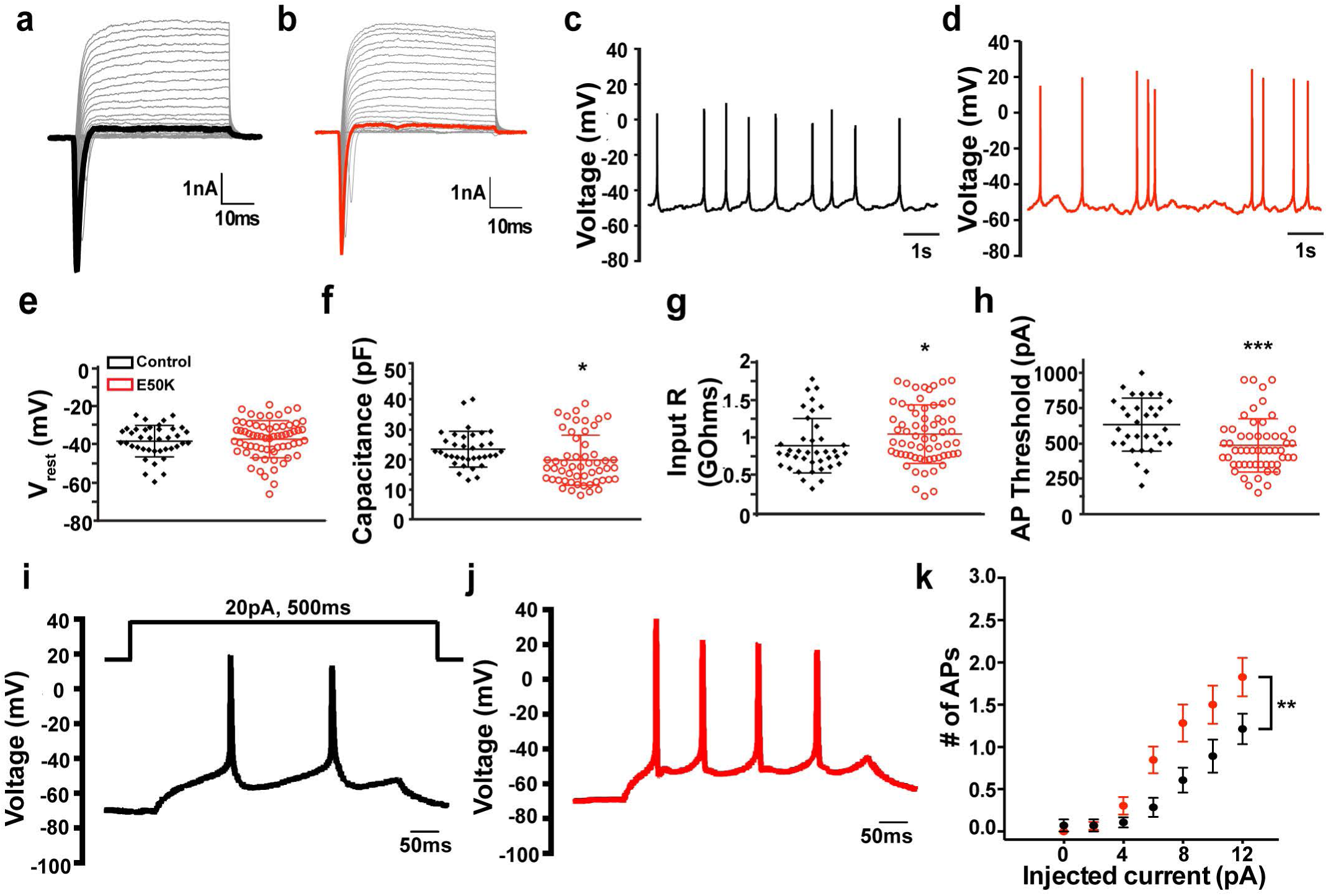
OPTN(E50K) RGCs display increased excitability. (a-d) OPTN(E50K) RGCs and isogenic controls displayed similarities in the ability to conduct inward sodium and outward potassium currents as well as the ability to fire spontaneous action potentials. (e) No significant differences were identified in the resting membrane potential between OPTN(E50K) RGCs and isogenic controls. (f-h) OPTN(E50K) RGCs demonstrated significantly lower cell capacitance, higher input resistance, and a lower action potential current threshold compared to isogenic controls. (i-k) Representative traces of evoked action potentials following current injections revealed the firing of significantly more action potentials in OPTN(E50K) RGCs compared to isogenic controls. Error bars represent SEM (n=2 biological replicates using H7 n=28 technical replicates and H7(E50K) hPSCs n=46 technical replicates) (*p < 0.05; **p < 0.01; ***p < 0.001).

### RNA sequencing demonstrates differential gene expression and pathways in OPTN(E50K) RGCs

To elucidate gene expression differences and the specific pathways affected by the OPTN(E50K) mutation, RNA sequencing was conducted on RGCs purified from OPTN(E50K) and isogenic control retinal organoids (Figure 5). Initial analysis demonstrated 75 downregulated genes and 117 upregulated genes associated with OPTN(E50K) RGCs when compared to isogenic control RGCs. Of the downregulated genes, a number of genes were associated with clearance of aggregated proteins, protein trafficking, and neurite outgrowth. Interestingly, genes found to be upregulated in OPTN(E50K) RGCs were related to retinal development as well as the OPTN gene. Pathway analysis revealed the upregulation and a variety of neurodegenerative pathways including Amyotrophic lateral sclerosis (ALS) and Alzheimer’s as well as a downregulation of pathways related to autophagy and neurite outgrowth. Thus, results from RNA sequencing demonstrated differential gene expression and pathways associated with OPTN(E50K) RGCs and allowed for further investigation into autophagy dysfunction as a result of the E50K mutation.

**Figure 5:**
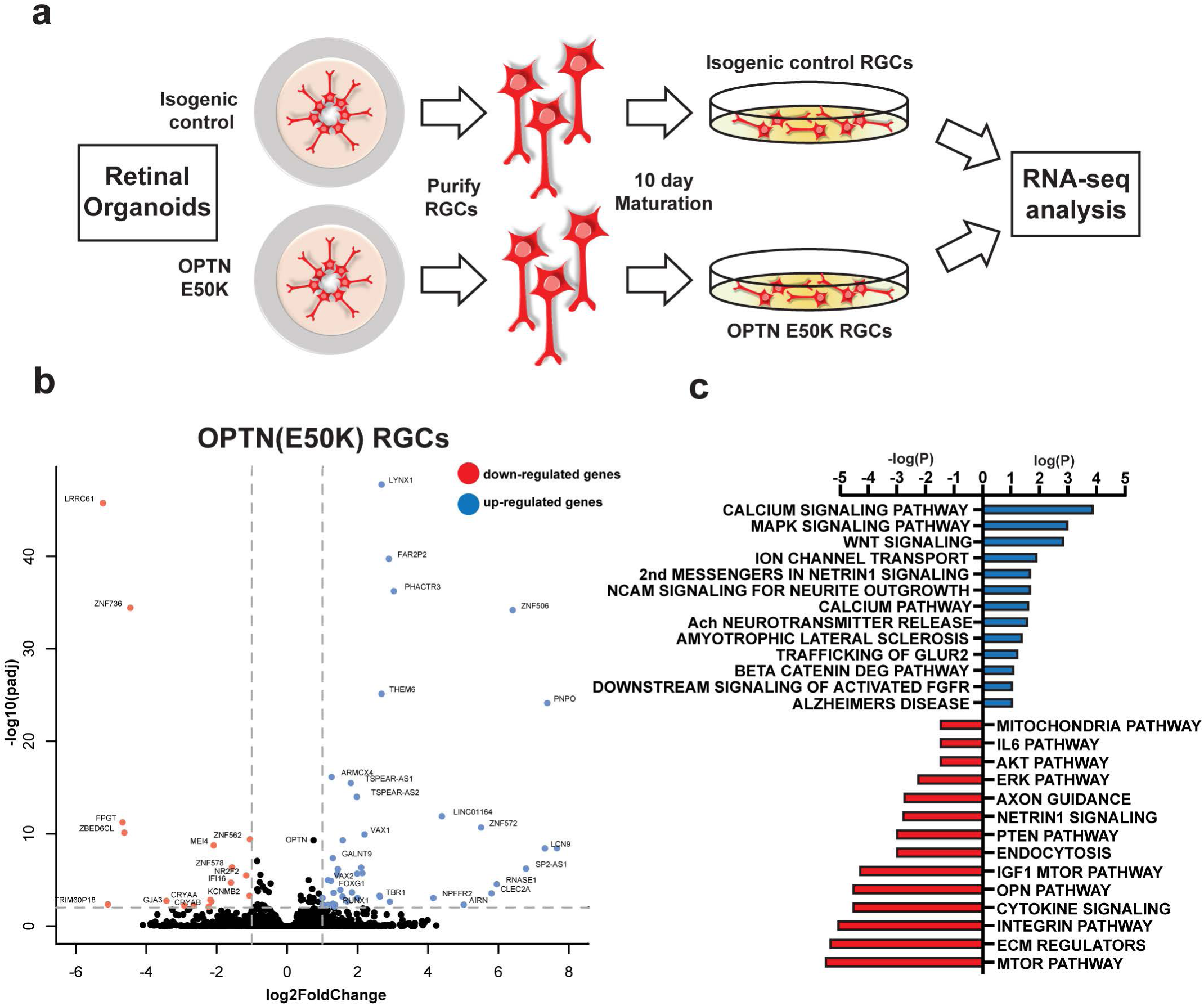
RNA sequencing revealed differential gene expression and pathway analysis in OPTN(E50K) RGCs. (a) A schematic demonstrates the work flow of RNA sequencing from OPTN(E50K) and isogenic control retinal organoids (n=4 for isogenic control and n=4 for OPTN(E50K). (b) Differential gene expression analysis demonstrated 75 down-regulated genes and 117 up-regulated genes in OPTN(E50K) RGCs when compared to isogenic control RGCs (p<0.05). (c) Pathway analysis revealed a variety of upregulated and downregulated pathways associated with glaucomatous neurodegeneration (p<0.1). n=4 biological replicates from both OPTN(E50K) hiPSCs and OPTN-corrected hiPSCs.

### OPTN(E50K) RGCs exhibit autophagy disruption and increased susceptibility to apoptosis

The OPTN protein is known to play an important role as an autophagy receptor, mediating the degradation of materials within the cell (Slowicka et al., 2016). Mutations to the OPTN protein, such as the E50K mutation, disrupt the autophagy pathway leading to the damage and degeneration of RGCs (Chalasani et al., 2014; Inagaki et al., 2018; Meng et al., 2012; Minegishi et al., 2013; Ying et al., 2015). As such, to elucidate possible disruptions to the autophagy pathway as a consequence of the OPTN(E50K) mutation in hPSC-derived RGCs, retinal organoids differentiated from both OPTN(E50K) and isogenic controls were examined for the expression of LC3AB, particularly the aggregation of this protein as a hallmark of autophagy dysfunction (Figure 6). Compared to isogenic controls, OPTN(E50K) organoids exhibited profound LC3AB aggregation exclusively within inner layers where RGCs reside, indicating a preferential effect upon these cells. To test the possibility of rescuing this phenotype, the autophagy pathway was enhanced through treatment with rapamycin, leading to a significant reduction of aggregates within inner layers of OPTN(E50K) organoids. As such, disruptions to the autophagy pathway could be a key contributor to the degeneration of OPTN(E50K) RGCs, with rescue of this phenotype by rapamycin suggesting the possibility of therapeutic intervention.

**Figure 6:**
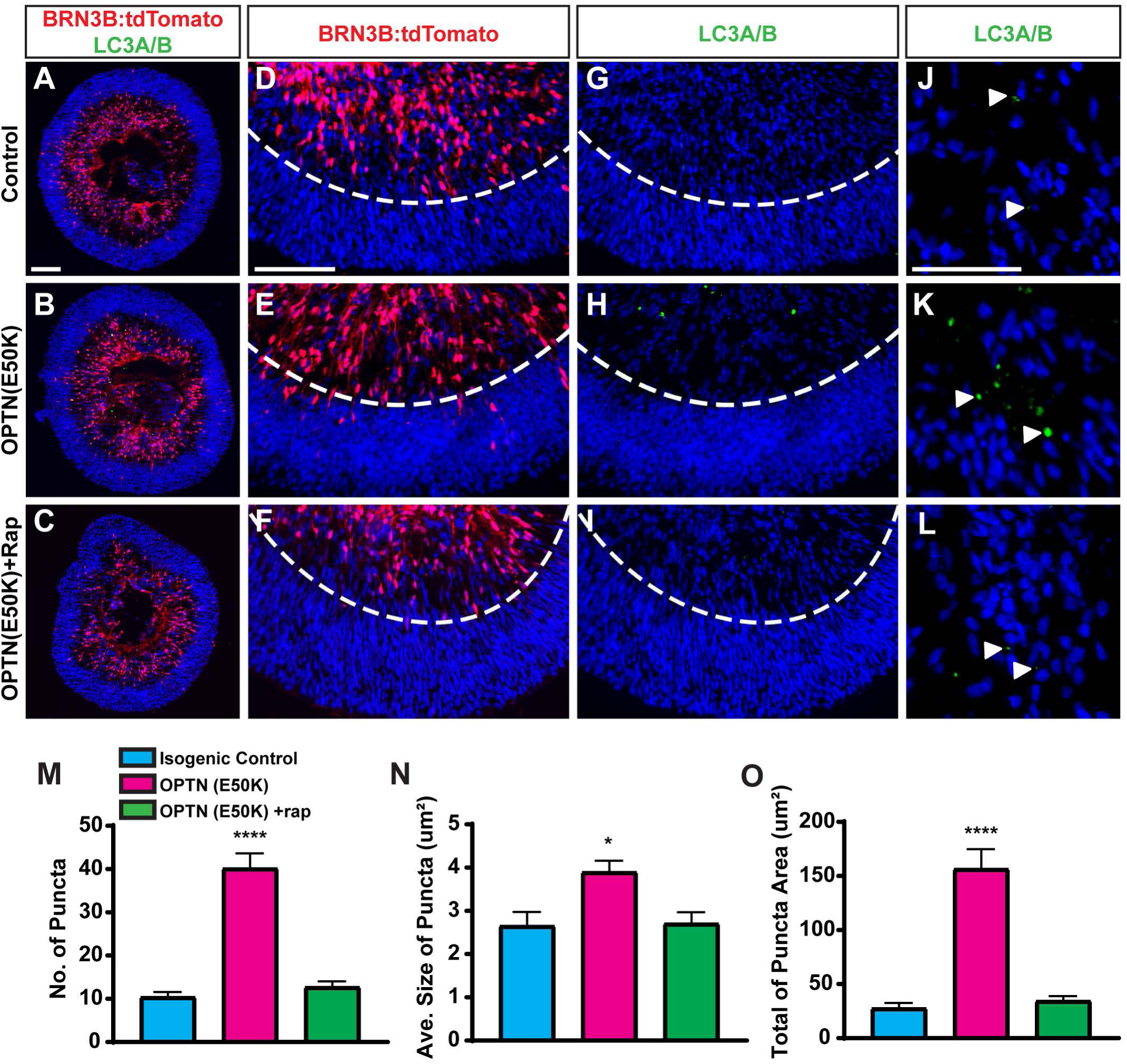
OPTN(E50K) retinal organoids demonstrate autophagy dysfunction through LC3A/B aggregation. (a-f) Retinal organoids from isogenic control, OPTN (E50K), and OPTN (E50K)+rapamycin conditions displayed proper organization, with BRN3B:tdTomato expression confined within inner layers. (g-l) LC3A/B aggregates were found specifically within the inner layers of OPTN(E50K) organoids compared to isogenic controls, with this aggregation rescued by rapamycin treatment. (m-o) Quantification of LC3AB aggregates demonstrated a significantly higher number of aggregates in OPTN(E50K) organoids compared to isogenic controls or rapamycin treated organoids, with these aggregates significantly larger in both puncta size and total area. Error bars represent SEM (n=3 biological replicates using OPTN(E50K) hiPSCs, OPTN-corrected hiPSCs, H7 and H7(E50K) hPSCs). (*p < 0.05; ****p < 0.0001). Scale bars are equal to 100 μm for a-i and 50μm for j-l.

To determine if autophagy disruption was correlated with a decrease in RGC viability, retinal organoids were similarly examined for apoptosis (Figure 7). Compared to isogenic controls, OPTN(E50K) organoids exhibited significantly more cleaved Caspase-3, with this increased expression found exclusively within inner layers of these organoids. Additionally, OPTN(E50K) organoids contained significantly fewer RGCs compared to isogenic controls based on the expression of BRN3, indicating a specificity of cell death to RGCs. Importantly, the number of photoreceptors were quantified by expression of OTX2, revealing no significant differences between OPTN(E50K) and isogenic control organoids (Supplemental Figure 3). Retinal organoids were then treated with rapamycin to determine if an association existed between apoptosis and autophagy. When treated with rapamycin, OPTN(E50K) organoids displayed decreased levels of cleaved Caspase-3, comparable with isogenic controls. As such, these results demonstrated an important link between autophagy dysfunction and apoptosis specifically within OPTN(E50K) RGCs.

**Figure 7:**
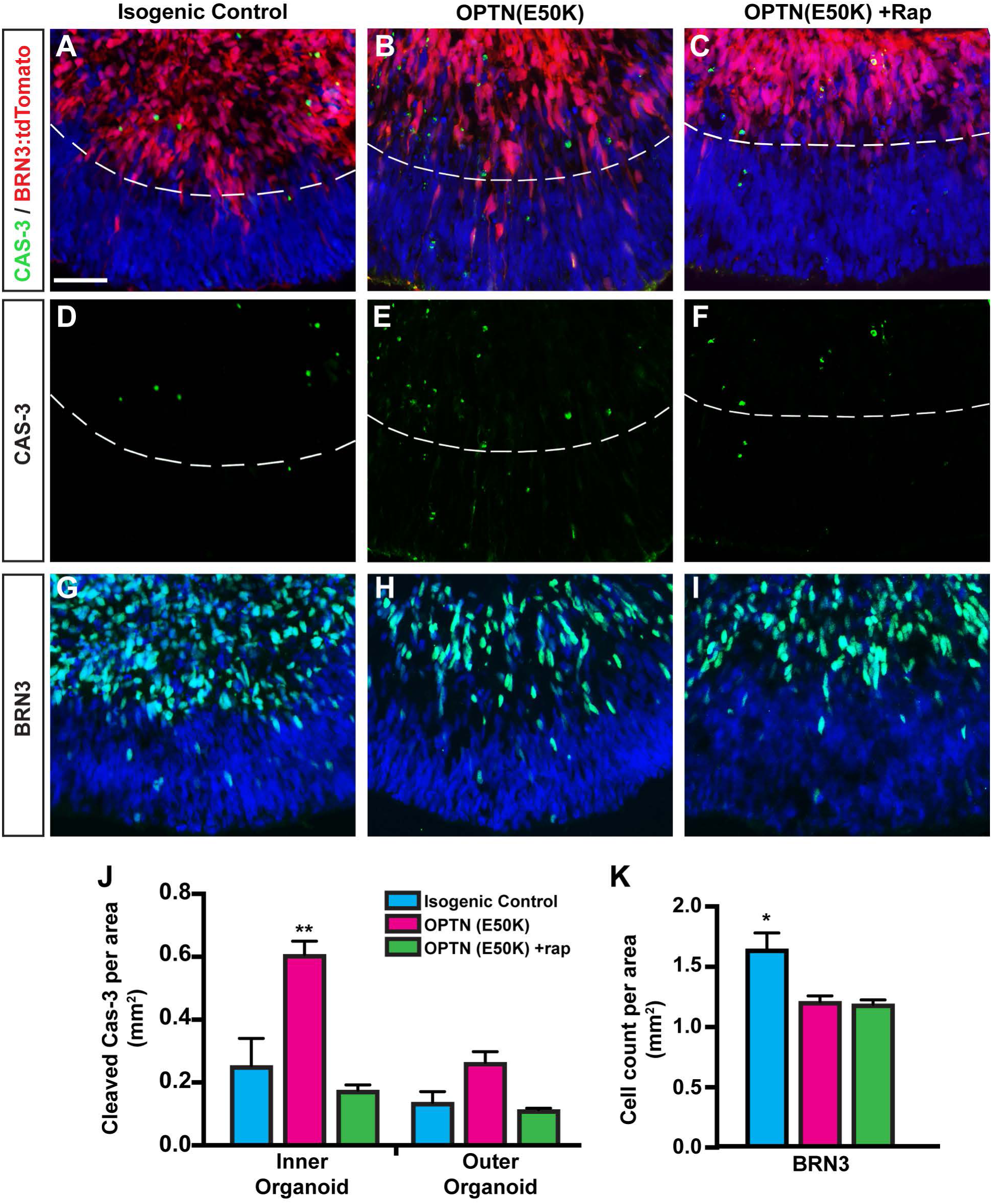
OPTN(E50K) retinal organoids display apoptotic activation and loss of RGCs. (a-f,j) OPTN(E50K) retinal organoids displayed significantly higher levels of cleaved CASPASE-3 within inner layers compared to outer layers as well as isogenic controls and OPTN(E50K) organoids treated with rapamycin. (g-i,k) OPTN(E50K) retinal organoids contained significantly lower numbers of RGCs based upon the expression of BRN3 compared to isogenic controls. Error bars represent SEM (n=3 biological replicates using OPTN(E50K) hiPSCs, OPTN-corrected hiPSCs, H7 and H7(E50K) hPSCs). (*p < 0.05; **p < 0.01). Scale bar equals 50μm.

## Discussion

hPSCs have been used as a reliable model system for studying both the development of many retinal cell types (Lamba et al., 2006; Meyer et al., 2011) as well as diseases which result in the degeneration of these cells (Sridhar, 2018). The current understanding of glaucomatous neurodegeneration has been established in part by the use of animal models mimicking RGC degeneration and injury (Agostinone et al., 2018; Kalesnykas et al., 2012; Tseng et al., 2015; Williams et al., 2013). Although these studies have been instrumental in discovering various disease phenotypes, important differences exist between rodent and primate retinas. As such, the use of hPSCs provides a powerful and complimentary model to study RGC development and disease that helps to bridge the gap between rodent studies and human glaucoma patients. Results of the current study demonstrate the most in depth phenotypic and functional characterization of RGCs grown from OPTN(E50K) hPSCs, revealing numerous neurodegenerative phenotypes including autophagy deficits and neurite retraction, allowing for a greater understanding of mechanisms leading to the degeneration and death of RGCs in glaucoma.

CRISPR/Cas9 gene editing provides new and exciting opportunities for the precise engineering of cells to generate new experimental models (Cong et al., 2013). Of particular interest for this study, CRISPR/Cas9 gene editing allows for the insertion of disease-causing mutations in hPSCs as well as the correction of these mutations in patient-specific hPSCs, minimizing effects due to variability between cell lines. As previous studies have demonstrated variability in the capacity of numerous hPSC lines to differentiate into retinal cells (Capowski et al., 2019; Meyer et al., 2011), this variability complicates the definitive identification of disease-related phenotypes apart from inherent differences between cell lines. The utilization of CRISPR/Cas9 gene editing in the current study provides the certainty required for the definitive identification of disease-related phenotypes directly linked to the OPTN(E50K) mutation. As such, the results of this study demonstrate the use of CRISPR/Cas9 technology for generation of hPSC-based RGC disease models, as well as their corresponding isogenic controls. Results of this study also provide the foundation for utilizing CRISPR/Cas9 gene editing of hPSCs for the study of other genetically inherited forms of glaucoma as well as other neurodegenerative diseases of the retina in order to discover novel mechanisms of degeneration and possible therapeutic targets.

The growth and differentiation of hPSCs as three-dimensional organoids provides an invaluable tool for the study of the retina, with retinal organoids following the spatio-temporal development of human retinogenesis (Nakano et al., 2012; Zhong et al., 2014). While previous studies have documented the use of retinal organoids to better understand RGC development within the retina (Fligor et al., 2018), the current study represents the use of retinal organoids as a tool for the study of RGC degenerative processes. In this capacity, several disease-related phenotypes were demonstrated to be specific to RGCs compared to other retinal cell types. However, as RGCs are the projection neurons of the retina that extend long and intricate neurites to transmit visual information to the brain, the morphological maturation and functionality of RGCs is sometimes easier to interpret when these cells are isolated from retinal organoids and grown in 2D culture (VanderWall et al., 2019). Results of the current study took advantage of both 3D organoid approaches as well as 2D RGC cultures to understand RGC development and maturation as well as identify disease phenotypes in these conditions.

The age-related loss of RGCs in glaucoma is characteristic of the loss of neurons found in many neurodegenerative diseases (Gupta and Yucel, 2007; Kovacs, 2017). In this context, RGCs differentiate normally during early retinogenesis, with degeneration associated with age and disease progression. As such, it is crucial for model systems of glaucoma to demonstrate this type of disease progression *in vitro*. Results of the current study indicated that OPTN(E50K)-hPSCs and isogenic controls differentiated into early optic cup-like retinal organoids that displayed a lamination of retinal layers. When organoids were grown in 2D cultures, no significant differences were observed in the expression of RGC-associated markers or the maturation in neuronal morphology at early stages. Disease phenotypes including neurite retraction, autophagy dysfunction, and apoptosis, were only identified in OPTN(E50K) cells after prolonged culture, with this progression recapitulating the age-related phenotypes observed in glaucomatous neurodegeneration.

The downregulation of RGC-associated transcription factors has been previously established in experimental glaucoma as well as RGC injury models, which provides an early indicator of degeneration in the retina (Soto et al., 2008; Weishaupt et al., 2005). Results of the current study demonstrated the downregulation of the RGC-associated transcription factors BRN3 and ISL1 in OPTN(E50K) RGCs compared to isogenic controls. Conversely, the expression of MAP2, which is also uniquely expressed by RGCs in the retina, was unchanged in OPTN(E50K) RGCs, providing a point of reference for those RGCs that have downregulated classical RGC-associated markers. Importantly, this downregulation of BRN3 and ISL1 was only identified after prolonged culture, with quantification at an earlier time points indicating no significant differences between OPTN(E50K) RGCs and isogenic controls. In future experiments using fluorescent reporters to identify RGCs in hPSC disease models, it will be necessary to account for the downregulation of these reporters to identify those RGCs at advanced stages of the disease state. It may also be necessary to develop reporter lines that retain their fluorescence even within the latest stages of the degenerative process.

The OPTN protein performs a variety of functions within the cell, including its role as an autophagy receptor (Slowicka et al., 2016). As such, OPTN interacts with a variety of essential autophagy proteins, with the E50K mutation in this protein leading to a disruption in this pathway. Autophagy pathway disruption has also been linked to neurodegeneration in a number of other diseases, including ALS, Alzheimer’s and Parkinson’s diseases, with the possibility of deficits in this pathway conserved as a mechanism leading to the degeneration of neurons (Chu, 2019). In the current study, results demonstrated profound aggregation of LC3AB in the presumptive RGC layers of OPTN(E50K) organoids compared to isogenic controls. Consequently, OPTN(E50K) organoids also exhibited significantly higher apoptosis within inner layers compared to both the outer layers of these same organoids as well as the presumptive RGC layers of isogenic controls. When treated with rapamycin, an activator of the autophagy pathway, aggregation of LC3AB and expression of cleaved Caspase-3 were reduced back to levels comparable to isogenic controls, reflecting an important balance between the autophagy and apoptotic pathways linked to the OPTN(E50K) mutation, with the potential to target these two pathways in the development of future therapeutic strategies.

The functional maturation of RGCs within the retina, including the firing of action potentials, is essential for the transmission of visual information from the retina to the brain (Wang et al., 1997), with changes in neuronal functionality associated with neurodegeneration. The overstimulation of cells through excitotoxic mechanisms has been implicated in a variety of neurodegenerative diseases including glaucoma (Cueva Vargas et al., 2015; Dzialo et al., 2013; Hynd et al., 2004). Results of the current study demonstrated that OPTN(E50K) RGCs exhibited a significantly lower action potential current threshold than isogenic controls, leading to the firing of significantly more action potentials, with these results suggesting the increased excitability of OPTN(E50K) RGCs as a key contributor to their degeneration. Importantly, both OPTN(E50K) and isogenic control RGCs were identified for patch clamp recordings through the expression of BRN3B:tdTomato fluorescence. Given that the results of this study also documented the downregulation of BRN3B:tdTomato in more advanced stages of OPTN(E50K) RGC maturation, the possibility exists that the phenotypes observed may represent an earlier stage of the degenerative process, with these phenotypes more severe in those RGCs which are more advanced in their degeneration and have downregulated the tdTomato reporter.

Although glaucoma is often overlooked as a neurodegenerative disease, it bears many similarities to other CNS diseases such as Parkinson’s, ALS, and Alzheimer’s which ultimately result in the degeneration of neurons in either the brain or spinal cord (Gupta and Yucel, 2007; Kovacs, 2017). Similar to previous studies of RGC damage as well as cortical neurons in Alzheimer’s disease (Agostinone et al., 2018; Agostinone and Di Polo, 2015; Dowjat et al., 1999; Kalesnykas et al., 2012; Williams et al., 2013), OPTN(E50K)-RGCs demonstrated significant neurite retraction and deficits in morphology compared to isogenic controls. The mechanisms to which RGC degeneration occurs in glaucoma remain somewhat inconclusive, with the most likely scenario including the interplay of numerous neurodegenerative mechanisms. Results found in this study identified numerous deficits including the downregulation of important RGC transcription factors, neurite retraction, autophagy disruption and apoptosis, as well as heightened excitability of RGCs, all of which have previously been associated with neurodegeneration. As such, the results and assays conducted in this study are not only relevant to the study of glaucoma, but also to many other diseases of the CNS, with the possibility to work collectively on targeted therapeutics at these distinctive pathways.

Overall, the results of the current study demonstrate a detailed characterization of the OPTN(E50K) mutation and how it affects the development and the degeneration of RGCs in a glaucomatous state. More so, this study extensively utilized CRISPR/Cas9 gene editing technology for the generation of disease models as well as isogenic controls for studies of glaucoma, with an important emphasis on the discrimination between mutation-causing phenotypes and cell line variability. In future studies using hPSC-based models of glaucomatous neurodegeneration, the identification of other unique neurodegenerative phenotypes and specific pathways leading to those phenotypes should be considered in order to better target therapeutic strategies.

## Methods

### CRISPR/Cas9 Gene Editing

The OPTN(E50K) mutation was inserted into H7BRN3B:tdTomatoThy1.2-hESCs (Thomson et al., 1998) (Sluch et al., 2017) and an hiPSC line harboring the OPTN(E50K) mutation (Ohlemacher et al., 2016) was corrected (See supplemental procedures for details on creation of gRNA, donor plasmids, and PCR primers). Before electroporation, hPSCs were acclimated to single cell passaging using Accutase. Electroporation was performed using the Neon transfection system (100 μl tip, Life Technologies) according to the manufacturer’s instructions. Briefly, 4 μg of pCas9-GFP and 6 μg of pUC57-HDR-U6-gRNA were mixed in 100 μl of buffer R. A pellet of 4 million hPSCs was resuspended in the mixture of buffer R and plasmids. Electroporation was performed under the following parameters: 1,500 V; 20 ms interval; 2 pulses. Subsequently, cells were plated onto Matrigel-coated plates in mTeSR1 medium with CloneR supplement (StemCell Technologies). 48 hours after electroporation, GFP positive cells were isolated by FACS to enrich for edited cells. After initial growth, clonal populations were isolated and expanded. To screen for the insertion/correction, genomic DNA from individual clones was extracted, and the portion of the OPTN gene containing the 50^th^ codon was amplified by PCR. This PCR product was then enzymatically digested by Hpy188III and run on a 1% gel. Properly edited clones were further confirmed by Sanger sequencing.

### Maintenance of hPSCs

hPSCs were cultured based on previously described protocols from 2 OPTN(E50K) diseased lines with corresponding isogenic controls as described in Figure 1 (Meyer et al., 2011; Ohlemacher et al., 2015). In brief, hPSC colonies were maintained on a matrigel-coated six well plate in mTeSR1 medium, with daily media changes. hPSCs were passaged every 4-5 days based upon their confluency. Prior to passaging, hPSCs were marked for areas of spontaneous differentiation and those areas were mechanically removed. At 70% confluency, hPSCs were enzymatically passaged with Dispase (2 mg/ml) for approximately 15 minutes and split at a ratio of 1:6 onto a new six well plate freshly coated with matrigel.

### Differentiation of RGCs

hPSCs were differentiated into a retinal organoids and RGCs using an established protocol with minor modifications (Meyer et al., 2011; Ohlemacher et al., 2015). Briefly, enzymatically passaged hPSC colonies were grown in suspension to produce embryoid bodies (EBs). Within the first three days of differentiation, EBs were slowly transitioned from mTesR1 medium into Neural Induction Media (NIM). At the sixth day of differentiation, BMP4 (50 ug/mL) was added to the EB flask containing NIM (Capowski et al., 2019). The same media was used at day 8 to induce adherence of EBs to a six well plate supplemented with 10% Fetal Bovine Serum. Half of the media was changed at day 9 and 12 to slowly reduce the concentration of BMP4 in the media. At day 15, cells were replenished with fresh NIM without BMP4 and the appearance of three-dimensional early optic vesicles was identified. The following day, early optic vesicle colonies were mechanically lifted and transferred into Retinal Differentiation Media, with a fresh media change every 2-3 days. At 30 days of differentiation, early retinal organoid cultures were supplemented with Glutamax and 2% FBS to aid in retinal organization. By day 45 of differentiation, retinal organoids were either enzymatically digested in Accutase to purify for RGCs, or preserved as floating organoid cultures for cryosectioning. For some experiments, retinal organoids were treated with rapamycin (2μM) for 24 hours and then fixed for immunocytochemistry.

To purify RGCs, retinal organoids were enzymatically dissociated into single cells using Accutase for 20 minutes at 37°C. Using previously described protocols, single cell suspensions were then enriched for RGCs with the Thy1.2 surface receptor using the MACS magnetic cell separation kit (Sluch et al., 2017). 10,000 RGCs were plated on poly-d-ornithine and laminin-coated 12 mm coverslips and maintained for up to 4 weeks in BrainPhys Media (VanderWall et al., 2019). To analyze neurite complexity, RGCs were transfected with GFP using lipofectamine 2 days before fixation to aid in identification of individual RGC neurites along with BRN3B:tdTomato expression.

### Immunocytochemistry

Samples were collected at selected time points within 2.5 months of differentiation. RGCs on coverslips and retinal organoids were fixed with 4% paraformaldehyde in phosphate buffer solution (PBS) for 30 minutes followed by 3 PBS washes. Retinal organoids were then prepared for cryosectioning through an incubation in 20% sucrose solution for 1 hour at room temperature, followed by an incubation in 30% sucrose solution overnight at 4°C. The following day, retinal organoids were transferred into OCT cryostat molds and placed on dry ice. 12μm thick sections were cut and used for immunocytochemical analyses.

Following fixation or sectioning, samples were permeabilized in 0.2% Triton X- 100 for 10 minutes at room temperature. Samples were then washed with PBS and blocked with 10% Donkey Serum for 1 hour at room temperature. Primary antibodies (Supplementary Table 1) were diluted in 0.1% Triton-X-100 and 5% Donkey Serum and applied overnight at 4°C. The next day, the primary antibody was removed and samples were washed 3 times with PBS and blocked with 10% Donkey Serum for 10 minutes at room temperature. Secondary antibodies were diluted at 1:1000 ratio in 0.1% Triton-X- 100 and 5% Donkey Serum and applied to samples for 1 hr at room temperature. The secondary antibodies were then removed and samples were washed 3 times with PBS before mounting onto slides for imaging. Immunofluorescent images were obtained using a Leica DM5500 fluorescence microscope.

### Quantification

Isogenic control and OPTN(E50K) RGCs were collected at 1, 2, 3, and 4 weeks after purification and analyzed based on neurite complexity, neurite length, and soma size. Several immunofluorescent images were taken of RGCs co-expressing tdTomato and GFP and soma area and neurite complexity were quantified using Image J and Photoshop, with the NeuronJ plugin used to quantify the length of RGC neurites.

Organoids from isogenic control, OPTN(E50K), and OPTN(E50K)+rapamycin sources were collected at 2.5 months of differentiation and quantified for expression of BRN3, OTX2, and CASPASE-3 using the cell counter plugin in Image J. LC3AB aggregate number, size, and area were quantified using the Image J particle analyzer. Organoid areas were also quantified using Image J plugins. For Caspase-3 quantification, organoid sections were imaged and the inner and outer layers were traced based on the expression of BRN3:tdTomato in Photoshop. The areas of inner and outer layers were determined using ImageJ and the cell counter plugin was used to quantify Caspase-3 fragments in each layer.

### Statistical analysis

Statistical significance for neurite complexity, soma size, neurite length, and electrophysiological recordings was performed using two-tailed student’s t test and significance based on a p value of less than 0.05. Significances for BRN3 and ISL1 quantification were achieved by a student’s t test based on a p value of less than 0.05. For BRN3, OTX2, CASPASE-3, and LC3 aggregates a One-Way Anova followed by a Tukey’s post hoc analysis was used to determine significance based on a p value less than 0.05.

### Electrophysiology

Whole-cell patch clamp recordings were performed at room temperature (∼22 °C) using a HEKA EPC-10 amplifier as previously described (VanderWall et al., 2019). The PatchMaster program was used for data acquisition. Electrodes were fabricated from 1.2mm borosilicate glass with a final resistance of 3-5MΩ. The extracellular solution contained (in mM): 140 NaCl, 1 MgCl_2_, 5 KCl, 2 CaCl_2_, 10 HEPES, 10 glucose and adjusted to a pH of 7.3 with NaOH. The pipette solution contained (in mM): 140 KCl, 5 MgCl_2_, 2.5 CaCl_2_, 5 EGTA, 10 HEPES and adjusted to a pH of 7.3 with KOH. RGCs were identified by tdTomato fluorescence. Resting membrane potential was measured with zero current injected and spontaneous firing was recorded if present. Input resistance was calculated according to V = IR with the change in voltage measured with a 200 ms, -100 pA stimulus. To enhance comparisons between cells during action potential activity recording, current was injected to bias the cell membrane potential to - 70 mV. Current threshold for action potential generation was obtained by a series of 1 ms stimuli of increasing intensity from 0 pA to 1100 pA in 50 pA steps from the biased membrane potential of -70 mV. The maximum number of action potentials elicited was measured either during a series of 500 ms stimuli of increasing intensity from 0 pA to 200 pA in 2pA or 10 pA steps from -70 mV or during a series of 1s ramp stimuli in 40pA increments. Voltage clamp recordings were obtained from each cell at the end of the series of current-clamp protocols. The peak amplitudes of sodium and potassium currents were measured using a standard I-V protocol with a holding potential at -80 mV. The current density was calculated by normalizing current amplitude to the capacitance of each cell.

### RNA sequencing prep and analysis

Organoids were immunopurified for RGCs after 45 days of differentiation and grown in adherent cultures for 10 days in BrainPhys medium. RNA was collected using the PicoPure RNA isolation kit. Total RNA was evaluated for its quantity and quality using an Agilent Bioanalyzer 2100. For RNA quality, a RIN number of 7 or higher was desired and 100ng of total RNA was used. cDNA library preparation included mRNA purification/enrichment, RNA fragmentation, cDNA synthesis, ligation of index adaptors, and amplification, following the KAPA mRNA Hyper Prep Kit Technical Data Sheet, KR1352 – v4.17 (Roche Corporate). Each indexed library was quantified and its quality accessed by Qubit and Agilent Bioanalyzer, and multiple libraries were pooled in equal molarity. The pooled libraries were denatured and neutralized before loading to NovaSeq 6000 sequencer at 300pM final concentration for 100b paired-end sequencing (Illumina, Inc.). Approximately 30M reads per library were generated. A Phred quality score (Q score) was used to measure the quality of sequencing and more than 90% of the sequencing reads reached Q30 (99.9% base call accuracy).

The sequencing data was next assessed using FastQC (Babraham Bioinfomatics, Cambridge, UK) and then mapped to the Human genome (GRCH38) using STAR RNA-seq aligner [23104886] with the parameter: “—outSAMmapqUNIQUE 60”. Uniquely mapped sequencing reads were assigned to GRCH38 reference genome using featureCounts. Genes with readcount per million (CPM) >0.5 in more than 3 of the samples were kept for further analysis. Differentially expressed genes were tested by using DESeq2 with FDR<0.05 as the significant cutoff [25516281]. Pathway enrichment analysis were conducted by hypergeometric test against Human Gene Ontology and MsigDB v6 canonical pathways, with p<0.01 as the significant cutoff [16199517]. For selected pathways, GSEA enrichment score and ssGSEA sample-wise enrichment score were used for visualization [16199517].

## Acknowledgements

We would like to thank Dr. Amelia Linneman for helpful discussions about autophagy analyses and the sharing of antibodies. We would also like to thank Dr. Don Zack and Dr. Valentin Sluch for sharing the BRN3B:tdTomato:Thy1.2 vectors used in the generation of RGC reporter cell lines used in the current study. Grant support was provided by the National Eye Institute (R01 EY024984 and R21 EY031120 to JSM), the Indiana Department of Health Spinal Cord and Brain Injury Research Fund (Grant #26343 JSM), and an Indiana CTSI Core Pilot grant (JSM). This publication was also made possible with partial support from a University Fellowship (KCH) and from an Indiana CTSI pre-doctoral research fellowship (UL1TR002529, A. Shekhar, PI) from the National Institutes of Health, National Center for Advancing Translational Sciences, Clinical and Translational Sciences Award (KBV).

## Author Contributions

KBV: experimental design, collection and analysis of data, manuscript writing. KCH: experimental design, collection and analysis of data, manuscript writing. YP: data collection and analysis. SL: data collection and analysis. CMF: data collection. AA: data collection. KL: data collection. PT: data collection and analysis. CZ: data collection and analysis. HCT: experimental design. TRC: experimental design and data analysis. JSM: experimental design, data analysis, manuscript writing, final approval of manuscript.

## References

Agostinone, J., Alarcon-Martinez, L., Gamlin, C., Yu, W.Q., Wong, R.O.L., and Di Polo, A. (2018). Insulin signalling promotes dendrite and synapse regeneration and restores circuit function after axonal injury. Brain : a journal of neurology 141, 1963–1980.

Agostinone, J., and Di Polo, A. (2015). Retinal ganglion cell dendrite pathology and synapse loss: Implications for glaucoma. Progress in brain research 220, 199–216.

Capowski, E.E., Samimi, K., Mayerl, S.J., Phillips, M.J., Pinilla, I., Howden, S.E., Saha, J., Jansen, A.D., Edwards, K.L., Jager, L.D., et al. (2019). Reproducibility and staging of 3D human retinal organoids across multiple pluripotent stem cell lines. Development (Cambridge, England) 146.

Chalasani, M.L., Kumari, A., Radha, V., and Swarup, G. (2014). E50K-OPTN-induced retinal cell death involves the Rab GTPase-activating protein, TBC1D17 mediated block in autophagy. PloS one 9, e95758.

Chu, C.T. (2019). Mechanisms of selective autophagy and mitophagy: Implications for neurodegenerative diseases. Neurobiology of disease 122, 23–34.

Cong, L., Ran, F.A., Cox, D., Lin, S., Barretto, R., Habib, N., Hsu, P.D., Wu, X., Jiang, W., Marraffini, L.A., et al. (2013). Multiplex genome engineering using CRISPR/Cas systems. Science (New York, NY) 339, 819–823.

Cueva Vargas, J.L., Osswald, I.K., Unsain, N., Aurousseau, M.R., Barker, P.A., Bowie, D., and Di Polo, A. (2015). Soluble Tumor Necrosis Factor Alpha Promotes Retinal Ganglion Cell Death in Glaucoma via Calcium-Permeable AMPA Receptor Activation. The Journal of neuroscience : the official journal of the Society for Neuroscience 35, 12088–12102.

Deng, F., Chen, M., Liu, Y., Hu, H., Xiong, Y., Xu, C., Liu, Y., Li, K., Zhuang, J., and Ge, J. (2016). Stage-specific differentiation of iPSCs toward retinal ganglion cell lineage. Molecular vision 22, 536–547.

Dowjat, W.K., Wisniewski, T., Efthimiopoulos, S., and Wisniewski, H.M. (1999). Inhibition of neurite outgrowth by familial Alzheimer’s disease-linked presenilin-1 mutations. Neuroscience letters 267, 141–144.

Dzialo, J., Tokarz-Deptula, B., and Deptula, W. (2013). Excitotoxicity and Wallerian degeneration as a processes related to cell death in nervous system. Archives italiennes de biologie 151, 67–75.

Fligor, C.M., Langer, K.B., Sridhar, A., Ren, Y., Shields, P.K., Edler, M.C., Ohlemacher, S.K., Sluch, V.M., Zack, D.J., Zhang, C., et al. (2018). Three-Dimensional Retinal Organoids Facilitate the Investigation of Retinal Ganglion Cell Development, Organization and Neurite Outgrowth from Human Pluripotent Stem Cells. Scientific reports in. press.

Gill, K.P., Hung, S.S., Sharov, A., Lo, C.Y., Needham, K., Lidgerwood, G.E., Jackson, S., Crombie, D.E., Nayagam, B.A., Cook, A.L., et al. (2016). Enriched retinal ganglion cells derived from human embryonic stem cells. Scientific reports 6, 30552.

Gupta, N., and Yucel, Y.H. (2007). Glaucoma as a neurodegenerative disease. Current opinion in ophthalmology 18, 110–114.

Hynd, M.R., Scott, H.L., and Dodd, P.R. (2004). Glutamate-mediated excitotoxicity and neurodegeneration in Alzheimer’s disease. Neurochemistry international 45, 583–595.

Inagaki, S., Kawase, K., Funato, M., Seki, J., Kawase, C., Ohuchi, K., Kameyama, T., Ando, S., Sato, A., Morozumi, W., et al. (2018). Effect of Timolol on Optineurin Aggregation in Transformed Induced Pluripotent Stem Cells Derived From Patient With Familial Glaucoma. Investigative ophthalmology & visual science 59, 2293–2304.

Kalesnykas, G., Oglesby, E.N., Zack, D.J., Cone, F.E., Steinhart, M.R., Tian, J., Pease, M.E., and Quigley, H.A. (2012). Retinal ganglion cell morphology after optic nerve crush and experimental glaucoma. Investigative ophthalmology & visual science 53, 3847–3857.

Kovacs, G.G. (2017). Concepts and classification of neurodegenerative diseases. Handbook of clinical neurology 145, 301–307.

Lamba, D.A., Karl, M.O., Ware, C.B., and Reh, T.A. (2006). Efficient generation of retinal progenitor cells from human embryonic stem cells. Proceedings of the National Academy of Sciences of the United States of America 103, 12769–12774.

Langer, K.B., Ohlemacher, S.K., Phillips, M.J., Fligor, C.M., Jiang, P., Gamm, D.M., and Meyer, J.S. (2018). Retinal Ganglion Cell Diversity and Subtype Specification from Human Pluripotent Stem Cells. Stem cell reports 10, 1282–1293.

Meng, Q., Lv, J., Ge, H., Zhang, L., Xue, F., Zhu, Y., and Liu, P. (2012). Overexpressed mutant optineurin(E50K) induces retinal ganglion cells apoptosis via the mitochondrial pathway. Molecular biology reports 39, 5867–5873.

Meyer, J.S., Howden, S.E., Wallace, K.A., Verhoeven, A.D., Wright, L.S., Capowski, E.E., Pinilla, I., Martin, J.M., Tian, S., Stewart, R., et al. (2011). Optic vesicle-like structures derived from human pluripotent stem cells facilitate a customized approach to retinal disease treatment. Stem cells (Dayton, Ohio) 29, 1206–1218.

Minegishi, Y., Iejima, D., Kobayashi, H., Chi, Z.L., Kawase, K., Yamamoto, T., Seki, T., Yuasa, S., Fukuda, K., and Iwata, T. (2013). Enhanced optineurin E50K-TBK1 interaction evokes protein insolubility and initiates familial primary open-angle glaucoma. Human molecular genetics 22, 3559–3567.

Minegishi, Y., Nakayama, M., Iejima, D., Kawase, K., and Iwata, T. (2016). Significance of optineurin mutations in glaucoma and other diseases. Progress in retinal and eye research 55, 149–181.

Nakano, T., Ando, S., Takata, N., Kawada, M., Muguruma, K., Sekiguchi, K., Saito, K., Yonemura, S., Eiraku, M., and Sasai, Y. (2012). Self-formation of optic cups and storable stratified neural retina from human ESCs. Cell stem cell 10, 771–785.

Ohlemacher, S.K., Iglesias, C.L., Sridhar, A., Gamm, D.M., and Meyer, J.S. (2015). Generation of highly enriched populations of optic vesicle-like retinal cells from human pluripotent stem cells. Current protocols in stem cell biology 32, 1h.8.1-20.

Ohlemacher, S.K., Sridhar, A., Xiao, Y., Hochstetler, A.E., Sarfarazi, M., Cummins, T.R., and Meyer, J.S. (2016). Stepwise Differentiation of Retinal Ganglion Cells from Human Pluripotent Stem Cells Enables Analysis of Glaucomatous Neurodegeneration. Stem cells (Dayton, Ohio) 34, 1553–1562.

Peng, Y.R., Shekhar, K., Yan, W., Herrmann, D., Sappington, A., Bryman, G.S., van Zyl, T., Do, M.T.H., Regev, A., and Sanes, J.R. (2019). Molecular Classification and Comparative Taxonomics of Foveal and Peripheral Cells in Primate Retina. Cell 176, 1222–1237.e1222.

Quigley, H.A. (2011). Glaucoma. Lancet (London, England) 377, 1367–1377.

Riazifar, H., Jia, Y., Chen, J., Lynch, G., and Huang, T. (2014). Chemically induced specification of retinal ganglion cells from human embryonic and induced pluripotent stem cells. Stem cells translational medicine 3, 424–432.

Slowicka, K., Vereecke, L., and van Loo, G. (2016). Cellular Functions of Optineurin in Health and Disease. Trends in immunology 37, 621–633.

Sluch, V.M., Chamling, X., Liu, M.M., Berlinicke, C.A., Cheng, J., Mitchell, K.L., Welsbie, D.S., and Zack, D.J. (2017). Enhanced Stem Cell Differentiation and Immunopurification of Genome Engineered Human Retinal Ganglion Cells. Stem cells translational medicine 6, 1972–1986.

Soto, I., Oglesby, E., Buckingham, B.P., Son, J.L., Roberson, E.D., Steele, M.R., Inman, D.M., Vetter, M.L., Horner, P.J., and Marsh-Armstrong, N. (2008). Retinal ganglion cells downregulate gene expression and lose their axons within the optic nerve head in a mouse glaucoma model. The Journal of neuroscience : the official journal of the Society for Neuroscience 28, 548–561.

Sridhar, A., Langer, K.B., Fligor C.M., Steinhart M., Miller, C.A., Ho-A-Lim, K.T., Ohlemacher S.K., Meyer, J.S. (2018). Human Pluripotent Stem Cells as In Vitro Models for Retinal Development and Disease. Regenerative Medicine and Stem Cell Therapty for the Eye, 17–49.

Tanaka, T., Yokoi, T., Tamalu, F., Watanabe, S., Nishina, S., and Azuma, N. (2015). Generation of retinal ganglion cells with functional axons from human induced pluripotent stem cells. Scientific reports 5, 8344.

Teotia, P., Chopra, D.A., Dravid, S.M., Van Hook, M.J., Qiu, F., Morrison, J., Rizzino, A., and Ahmad, I. (2017a). Generation of Functional Human Retinal Ganglion Cells with Target Specificity from Pluripotent Stem Cells by Chemically Defined Recapitulation of Developmental Mechanism. Stem cells (Dayton, Ohio) 35, 572–585.

Teotia, P., Van Hook, M.J., Wichman, C.S., Allingham, R.R., Hauser, M.A., and Ahmad, I. (2017b). Modeling Glaucoma: Retinal Ganglion Cells Generated from Induced Pluripotent Stem Cells of Patients with SIX6 Risk Allele Show Developmental Abnormalities. Stem cells (Dayton, Ohio) 35, 2239–2252.

Thomson, J.A., Itskovitz-Eldor, J., Shapiro, S.S., Waknitz, M.A., Swiergiel, J.J., Marshall, V.S., and Jones, J.M. (1998). Embryonic stem cell lines derived from human blastocysts. Science (New York, NY) 282, 1145–1147.

Tseng, H.C., Riday, T.T., McKee, C., Braine, C.E., Bomze, H., Barak, I., Marean-Reardon, C., John, S.W., Philpot, B.D., and Ehlers, M.D. (2015). Visual impairment in an optineurin mouse model of primary open-angle glaucoma. Neurobiology of aging 36, 2201–2212.

VanderWall, K.B., Vij, R., Ohlemacher, S.K., Sridhar, A., Fligor, C.M., Feder, E.M., Edler, M.C., Baucum, A.J., 2nd, Cummins, T.R., and Meyer, J.S. (2019). Astrocytes Regulate the Development and Maturation of Retinal Ganglion Cells Derived from Human Pluripotent Stem Cells. Stem cell reports 12, 201–212.

Wang, G.Y., Ratto, G., Bisti, S., and Chalupa, L.M. (1997). Functional development of intrinsic properties in ganglion cells of the mammalian retina. Journal of neurophysiology 78, 2895–2903.

Weishaupt, J.H., Klocker, N., and Bahr, M. (2005). Axotomy-induced early down-regulation of POU-IV class transcription factors Brn-3a and Brn-3b in retinal ganglion cells. Journal of molecular neuroscience : MN 26, 17–25.

Williams, P.A., Howell, G.R., Barbay, J.M., Braine, C.E., Sousa, G.L., John, S.W., and Morgan, J.E. (2013). Retinal ganglion cell dendritic atrophy in DBA/2J glaucoma. PloS one 8, e72282.

Ying, H., Turturro, S., Nguyen, T., Shen, X., Zelkha, R., Johnson, E.C., Morrison, J.C., and Yue, B.Y. (2015). Induction of autophagy in rats upon overexpression of wild-type and mutant optineurin gene. BMC cell biology 16, 14.

Zhong, X., Gutierrez, C., Xue, T., Hampton, C., Vergara, M.N., Cao, L.H., Peters, A., Park, T.S., Zambidis, E.T., Meyer, J.S., et al. (2014). Generation of three-dimensional retinal tissue with functional photoreceptors from human iPSCs. Nature communications 5, 4047.

