## Supplemental Info for "Retinal ganglion cells harboring the OPTN(E50K) mutation exhibit neurodegenerative phenotypes when derived from hPSC-derived three dimensional retinal organoids"

**Supplemental Table 1:** Primary Antibody Inventory

| Primary Antibody | Company | Catalog Number | Dilution |
|---|---|---|---|
| BRN3-pan | Santa Cruz | Sc-514474 | 1:200 |
| Cleaved Caspase-3 | Promega | G7481 | 1:200 |
| Chx10 | Santa Cruz | Sc-21690 | 1:200 |
| Histone H3 | Cell Signaling | 9715S | 1:1000 |
| ISLET1 | DSHB | 40.2D6 | 1:100 |
| LC3AB | Cell Signaling | 12741 | 1:100 |
| MAP2 | Santa Cruz | Sc-19344 | 1:200 |
| Nanog | Stemgent | 09-0020 | 1:100 |
| Neurexin-1 | Millipore | Abn161-I | 1:1000 |
| Otx2 | R&D Systems | Af1979 | 1:2000 |
| Pax6 | DSHB | Pax6 | 1:50 |
| PSD-95 | Abcam | Ab18258 | 1:500 |
| RBPMS | PhosphoSolutions | 1830-RBPMS | 1:500 |
| Recoverin | Millipore | Ab5585 | 1:2000 |
| RFP | Rockland | 600-401-379 | 1:1500 |
| SNAP-25 | Millipore | Mab331 | 1:500 |
| SNCG | Abcam | Ab55424 | 1:100 |
| Tra1-81 | Stemgent | 09-0011 | 1:1000 |

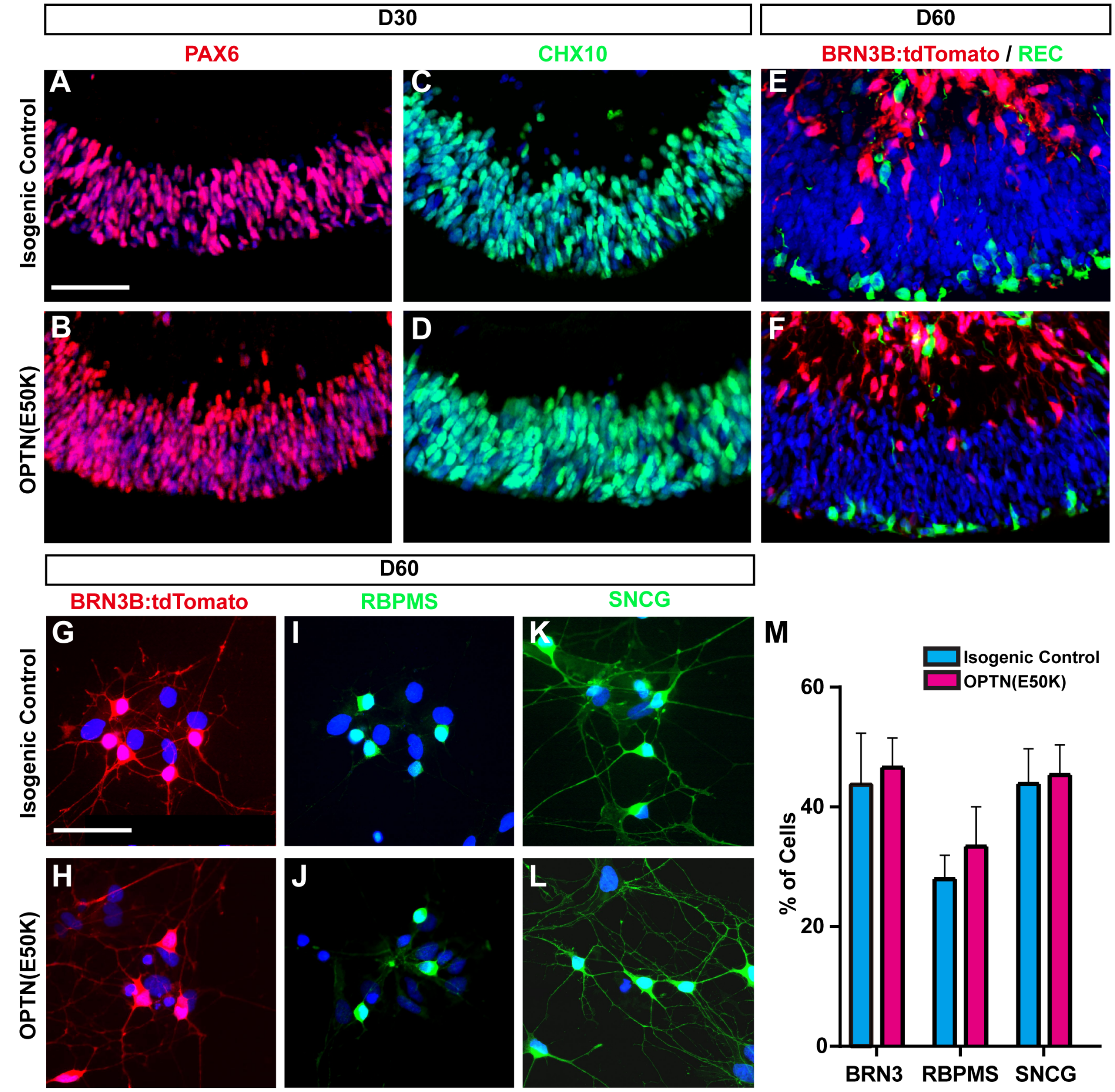


**Supplemental Figure 1. OPTN(E50K) and isogenic controls differentiate similarly in early developmental stages.** (a-d) ICC expression of PAX6 and CHX10 at day 30 revealed no differences between OPTN(E50K) and isogenic control retinal organoids. (e-f) Similarly, both conditions demonstrated similar expression patterns for BRN3B:tdTomato within inner layers and Recoverin within outer layers of retinal organoids after 60 days of differentiation. (g-m) RGCs purified from OPTN(E50K) and isogenic control retinal organoids demonstrated no significant differences in the expression of BRN3B:tdTomato, RBPMS, and SNCG. Error bars represent SEM. Scale bars equal 100μm for a-f and 50μm for g-l.

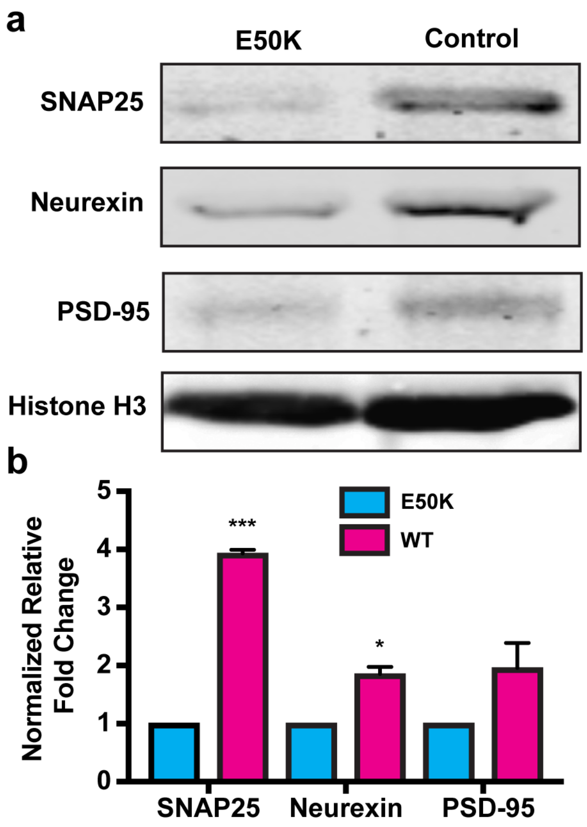


**Supplemental Figure 2**: **OPTN(E50K) RGCs demonstrate decreased synaptic proteins.** (a) Western blot analysis revealed significantly decreased synaptic proteins in OPTN(E50K) RGCs compared to isogenic controls. (b) Normalized densitometry indicated a significant difference in the amount of protein between OPTN(E50K) RGCs and isogenic controls. Error bars equal SEM.

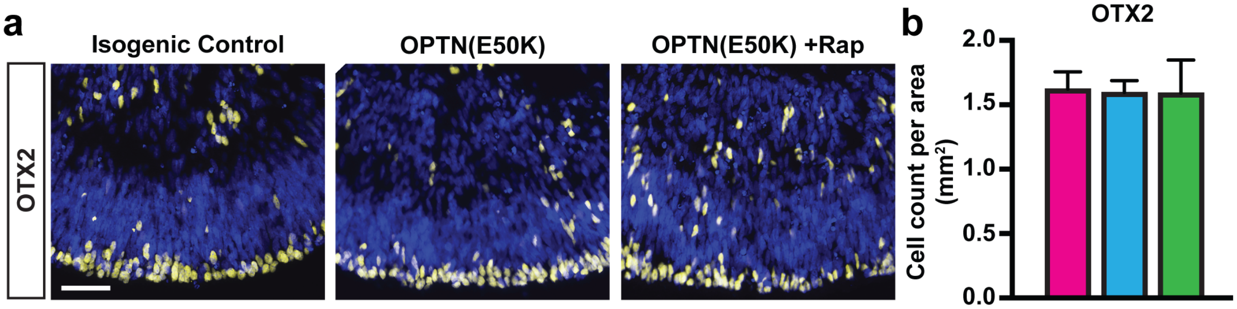


**Supplemental Figure 3: OPTN(E50K) and isogenic control organoids express a similar number of photoreceptors.** (a-b) Immunostaining demonstrated similar expression in the number of OTX2-expressing photoreceptors. (b) Quantification revealed no significant differences in the number of OTX2 cells between each of the conditions.

**Supplemental Methods:**

**Western Blot**

Samples were collected in 2% SDS solution within 4 weeks of plating. 4x sample buffer +100uM DTT was added to cell lystates and incubated at 70$^{o}$C for 10 minutes. Samples were then loaded into a 4-15% gradient pre-cast gel. The Trans-Blot Turbo system (BioRad) was utilized to transfer the completed gel onto nitrocellulose and a ponso stain was performed to determine total protein. Following transfer, the nitrocellulose was blocked using 5% milk in tris buffered saline (TBS) plus 0.1% Tween-20 (T) for 20 minutes and then incubated with primary antibodies overnight at 4$^{o}$C. The following day, the blot was washed 3x with 5% milk in TBS-T and the appropriate secondary antibody was applied for 1 hour at room temperature in the dark. Following secondary antibody incubation, blots were washed 3x with TBS and imaged using the Li-COR Odyssey CLx imaging system. Fluorescence intensities were calculated for each band and normalized to a loading control.
